# Honey bee drones are synchronously hyperactive inside the nest

**DOI:** 10.1101/2023.01.19.524638

**Authors:** Louisa C. Neubauer, Jacob D. Davidson, Benjamin Wild, David M. Dormagen, Tim Landgraf, Iain D. Couzin, Michael L. Smith

**Author notes:** Corresponding author: Michael L. Smith. These authors contributed equally to this work (Louisa C. Neubauer, Jacob D. Davidson).

## Abstract

Eusocial insects operate as an integrated collective with tasks allocated among individuals. This applies also to reproduction, through coordinated mating flights between male and female reproductives. While in some species male sexuals take only a single mating flight and never return, in the Western honey bee, *Apis mellifera*, the male sexuals (drones) live in the colony throughout their lives. Prior research has focused almost exclusively on drone behavior outside of the nest (mating flights), while ignoring the majority of their life, which is spent inside the nest. To understand the in-nest behavior of drones across their lives, we used the BeesBook tracking system to track 192 individually-marked drones continuously for over 20 days, to examine how drones moved and spent time in the nest. In agreement with previous work, we find that drones spend most of their time immobile at the nest periphery. However, we also observe that drones have periods of in-nest hyperactivity, during which they become the most active individuals in the entire colony. This in-nest hyperactivity develops in drones after age 7 days, occurs daily in the afternoon, and coincides with drones taking outside trips. We find strong synchronization across the drones in the start/end of activity, such that the drones in the colony exhibit a “shared activation period”. The duration of the shared activation period depends on the weather; when conditions are suitable for mating flights, the activation period is extended. At the individual-level, we see that the activation order changes from day to day, suggesting that both the external influence of weather conditions, as well as exchange of social information, influences individual activation. Using an accumulation-to-threshold model of drone activation, we show that simulations using social information match experimental observations. These results provide new insight into the in-nest behavior of drones, and how their behavior reflects their role as the male gametes of the colony.

## 2 Introduction

The parts of an organism can be defined by their functional role (e.g. gas-exchange, digestion, locomotion) or their ultimate role: soma versus gametes (Weismann, 1893). This same concept applies to eusocial insect colonies – workers can be organized into functional subgroups that carry out tasks, similar to organs, but colonies can also be defined according to those that form the soma (workers) versus gametes (sexuals) (Seeley, 1989; Smith and Szathmáry, 1995; Beshers and Fewell, 2001; Hölldobler et al., 2009; O’Shea-Wheller et al., 2021). Male reproductives in honey bee colonies, drones, are the equivalent of colony-level sperm, whose goal is to mate with virgin queens from other colonies (Ruttner and Ruttner, 1966; Loper et al., 1992; Koeniger et al., 2005; Woodgate et al., 2021). Drones are important for a colony’s reproductive success, and each virgin queen mates with multiple drones to obtain sperm from diverse patrilines – a critical contribution for colony-level function (Tarpy, 2003; Jones et al., 2004; Mattila and Seeley, 2007; Mattila et al., 2012).

In many social insect systems, sexuals are reared and depart only on a single mating flight, never to return (e.g. *Solenopsis invicta* (Tschinkel, 2006); *Reticulitermes speratus* and *Coptotermes formosanus* (Mizumoto and Dobata, 2019)). Across the bees, male mating strategies depend on patterns of female emergence, nest distribution, and male density (Paxton, 2005). The behavioral outcomes of these mating strategies, however, focus on male behavior outside of the nest, while neglecting their in-nest behavior. In the honey bees, genus Apis, drones live their entire lives in/on the nest, departing on multiple mating flights per day (Oldroyd and Wongsiri, 2006; Seeley, 2019). Therefore, even if their ultimate purpose is solely related to colony reproduction, the fact that they depart and repeatedly return shows that drones are part of the colony’s overall organization within the nest. Previous research has examined both the in- and out-of-nest behavior of workers – who perform the bulk of colony functions (Seeley, 1995; Hölldobler et al., 2009; Seeley, 2010) – but prior work on drones has focused almost exclusively on their behavior outside the nest, specifically their mating flights.

Drone mating flights occur across all species of *Apis* (Oldroyd and Wongsiri, 2*006). Each Apis* species has its own time-window for drone departures, which helps reproductive isolation in regions where multiple species of *Apis* coincide (Koeniger et al., 1988; Won, 1996; Oldroyd and Wongsiri, 2006). In the Western honey bee, *A. mellifera*, drones depart on multiple mating flights each afternoon, during which they visit multiple drone congregation areas (Oertel, 1956; Taber, 1964; Ruttner, 1966; Reyes et al., 2019; Woodgate et al., 2021). Drones that successfully mate will die, but there are thousands of drones produced for each available virgin queen (Smith et al., 2016), so most drones depart on daily flights without ever mating. Flight activity depends on the drone’s maturation; young drones perform short orientation flights (Oertel, 1956; Witherell, 1971), and older drones depart on long-distance mating flights 1-5 km from the nest (Ruttner and Ruttner, 1972; Reyes et al., 2019). Previous work has shown that drone flights tend to occur only under specific weather conditions – temperature above 20^*◦*^C, light intensity around 2000 Lux, no cloud cover, and wind speed below 30 km/h (Koeniger, 1986; Neves et al., 2011; Reyes et al., 2019). These environmental conditions align with the conditions needed for virgin queens to depart on mating flights (Taber, 1964; Koeniger, 1986). Indeed, virgin queens need to be particularly choosy about weather conditions – if they fail to return home, the colony is rendered “hopelessly queenless” and will perish (Smith, 2018). At the end of the summer, when there are no more virgin queens with whom to mate, drones are no longer useful to the colony, and workers evict them to die of starvation outside the nest (Morse et al., 1967; Free and Williams, 1975; Wharton et al., 2008; Cicciarelli, 2013).

These details describe the outside activities of drones, but less is known about their behavior within the nest. Previous work found that drones tend to remain stationary on the combs, only moving to feed (Free et al., 1957). Young drones tend to be located in the center brood-rearing region of the nest, whereas older drones are located at the nest periphery (Fukuda and Ohtani, 1977). Drones do not perform colony work, though drones will contribute to nest thermoregulation by heating themselves when cold (Kovac et al., 2009).

Here, we examine the in-nest behavior of drones of the Western honey bee, by tracking the movement patterns and spatial positioning of individual drones in an observation hive over their entire lives. We find that drones exhibit a daily period of in-nest hyperactivity, during which they also take trips outside of the nest. The duration and onset of the high-activity period depends on weather conditions. Looking at individual drones, we find that the start and end of the activation period is highly synchronized, and that individual drone activation order is not consistent from day to day. Finally, we formulate an accumulation-to-threshold model of drone activation, and examine simulation results with and without social information, in comparison to experimental observations. These results provide new insight into the in-nest behavior of drones as the male reproductive units of the colony.

## 3 Methods

### 3.1 Observation hive and recording

This study was performed at the University of Konstanz, Germany (47.6894N, 9.1869E). On 10 May 2019, the observation hive was installed with a single queen, 4000 unmarked workers, and three frames containing brood and honey (observation hive dimensions: 490×742 mm; ‘Deutsche-Normal’ frames: 395×225 mm). From June 5 to September 23, 2019, individually-marked newborn workers were introduced to the observation hive colony every 4 to 5 days (250-400 newborns per introduction). On June 7 and 12, when colonies were producing drones, we individually marked and introduced drones to the same observation hive colony (192 total: 160 and 32, respectively). Newborn bees were sourced from colonies headed by naturally mated queens from the University of Konstanz apiary (*Apis mellifera carnica*). All newborns were hatched overnight in an incubator, so the size of the cohort matches the number of individuals that would have naturally emerged overnight. The incubator was set to 34^*◦*^C and 50% RH, and newborns were marked that morning with individual BeesBook tags (Wario et al., 2015; Boenisch et al., 2018). BeesBook tags are printed on paper and attached to the thorax of bees, and remain attached for their whole lives.

From June 5 to October 23, 2019, the observation hive was recorded at 3 frames per second with 4 Basler acA4112-20um cameras fitted with Kowa LM25XC lenses. To mimic natural conditions inside a nest, the colony was illuminated with infrared light (850nm 3W LED’s), which the bees cannot perceive (Peitsch et al., 1992), and the colony had free access to the outside via a 2cm diameter entrance tunnel. This investigation focuses on the recording period during which drones were present in the colony, which began June 7, 2019.

To create a map of the nest, we periodically traced the contents of the observation hive onto plastic sheets by outlining the following: honey storage, pollen storage, brood, empty comb, wooden frames, peripheral galleries, and dances observed on the dance floor (as in Smith et al. (2016)). These plastic sheets were then scanned with an architectural scanner (Ruch-Medien, Konstanz), and digitized.

### 3.2 Data processing

Using the BeesBook tracking system, the raw images were processed to detect and decode individually marked bees (Wario et al., 2015; Wild et al., 2021). These processed data contain tag ID, ID detection confidence, position in the nest, bee orientation, and time. All data was stored in a PostreSQL database. The death date of each marked individual was estimated using a Bayesian changepoint model (as in Wild et al. (2021)). This method accounts for a low rate of erroneous detections in bees that have already died, and time periods when individuals are observed less frequently or not at all (e.g. while foraging). An individual’s death date was used as a cutoff for including data in subsequent calculations. Given that drones have a low probability of mating success per flight (thousands of drones produced per virgin queen (Smith et al., 2016)), and high potential for mortality (Visscher and Dukas, 1997), we did not attempt to differentiate drones that successfully mated and died versus those that simply died while outside the nest, or those that died within the nest of other causes.

We use the processed trajectory data to calculate metrics that represent the behavior of each individual. This is done by averaging over time bins of duration *T*, where *T* = 5 minutes was used for all analyses shown here. All data points used in the analysis were above a detection confidence threshold of 0.8 (detection confidence is an output of the BeesBook tracking system that indicates how likely a detection is correct - see (Boenisch et al., 2018)). We calculated behavioral metrics for each bee that had a minimum of 10 detections in a time bin. In general, this choice was made in order to keep as much data as possible during the data processing step, while also filtering out likely errors, and maintaining flexibility in possible analyses with the data. For the analysis of drone behavior we focus on speed and time spent outside; however, the full list of all calculated metrics include quantities to represent space use within the nest (time on honey, brood, or dance floor, and exit distance), detection (time observed, time outside, number of outside trips, and number of dance floor visits), and movement/spatial localization (speed, circadian coefficient, dispersion, fraction of nest visited) - see Smith et al. (2022) for full details on how these metrics are calculated. For each day, we calculated behavioral parameters for each bee, averaged in the bins of duration *T* (June 7 and June 12 cohorts: 736 workers and 192 drones, total). For all averages shown in the figures, we used a per-bee average of the calculated metrics.

To estimate when a bee was outside, we used an algorithm with input from the binned calculations for detection and exit distance. Although we do not have direct observations or detections corresponding to exiting the hive (i.e. there was not a camera at the hive exit that could read barcodes), leaving the nest means that there will be gaps in the detection of individual bees for some period. However, it is also possible that a bee’s barcode in the observation hive is not always detected, for example if the bee is upside-down or in a dense crowd of other bees. Because of this we use both detection and exit distance to estimate when bees were out of the nest. We first calculate the time observed and median exit distance in 5-minute bins over the course of a day. A bee is then estimated to have exited the nest in a time bin *t*_*exit*_ if the time observed in *t*_*exit*_ is less than a threshold of *t*_*obs*_=2 sec., and if the median exit distance in time bin *t*_*exit*_ *−* 1 is less than a threshold *d*_*exit*_=31.25 cm (2500 pixels). The bee is considered to have re-entered in bin *t*_*enter*_ if the time observed in *t*_*enter*_ is greater than or equal to *t*_*obs*_. The values *t*_*obs*_ and *d*_*exit*_ are analysis parameters, and the results can depend strongly on the choice of *d*_*exit*_; we choose the value of 31.25 cm to represent a feasible median exit distance for a bee traveling to the exit during a 5-minute period (see Figure S2 for exit distances labeled within the observation hive). With these results, we determine multiple instances of exit and re-entry times during the course of a day, and use this to calculate the number of outside trips (the number of times a bee is estimated to have exited the nest), as well as the time spent outside.

All data associated with this study is freely available online: see Smith et al. (2023).

## 4 Results

### 4.1 In-hive movement of drones over time

We individually tagged and introduced 192 newly-emerged drones into an observation hive, and tracked their motion continuously at 3 fps over 20+ days using the BeesBook automated tracking system (Wario et al., 2015; Boenisch et al., 2018; Wild et al., 2021). The drones were divided into two cohorts with birth dates 5 days apart. Two worker cohorts were introduced simultaneously with the drones (bees within each cohort have the same date of birth: worker/drone cohort 1, June 7; worker/drone cohort 2, June 12; mid-June is when colonies are naturally producing drones). We quantify the individual behavior of drones by calculating the median speed of individuals in 5-minute bins, as well as using position and detection within the observation hive to estimate when individuals leave the nest (see Methods). We use nest tracing to map the comb contents in the observation hive (honey stores, brood care, pollen stores, festoon for comb-building; as in Smith et al. (2016, 2022)), so that the position of bees in the hive can be compared with current nest structure and use.

In agreement with previous observations (Free et al., 1957), we found that drones remained immobile in the nest throughout most of the day (Per-drone median speed during the observation period: 0.08 ± 0.027 cm/sec. Per-worker speed for workers in cohorts 1&2 during this period: 0.262 ± 0.035 cm/sec. Two-sample t-test, *p <* 0.001). However, we also found that drones had extreme bouts of hyperactivity (Figure 1A). During these hyperactive bouts, drones’ speed was several times faster than workers (Per-drone median speed during shared activation periods: 0.779 ± 0.282 cm/sec; Per-worker speed for workers in cohorts 1&2 during shared activation periods: 0.355 ± 0.068. Two-sample t-test, *p <* 0.001; see below, and Methods, for definition of shared activation period). This demonstrates that while drones do spend the majority of their time immobile, or barely moving, they can also be the fastest individuals in the colony, albeit during a limited time-window (see trajectories in Supplementary Video 1).

**Figure 1:**
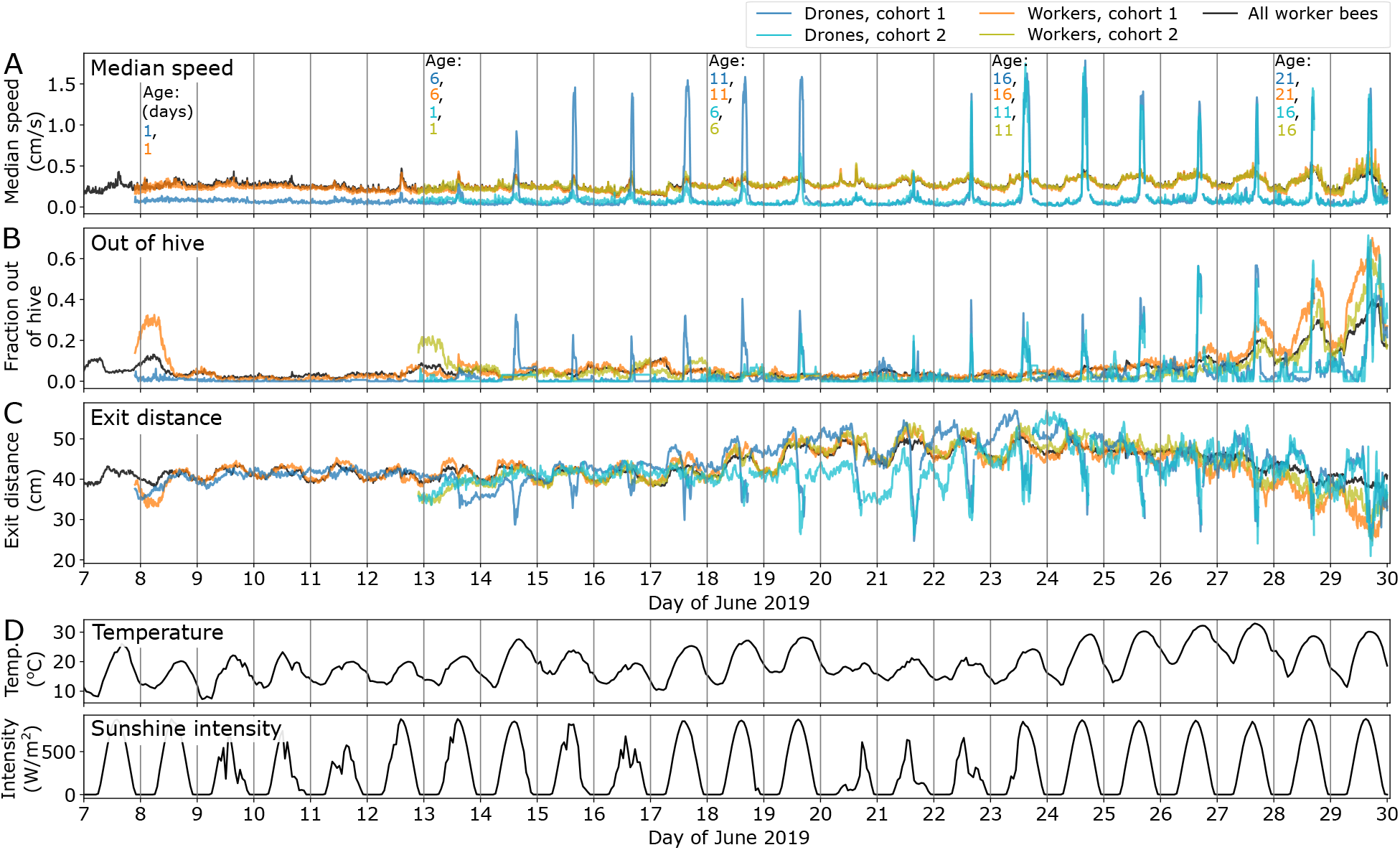
Movement of drones and workers, and weather over time. (A) Median speed, (B) Fraction of bees out of the hive, and (C) Median exit distance for each drone and worker cohort, over the observation period. Drone and worker cohorts 1 were introduced on June 7; drone and worker cohorts 2 on June 12. Each line shows the per-bee average across the group, calculated using time bins of 5 minutes (see Methods for data processing details). The black line shows values for all marked worker bees in the observation hive (this includes 3246 bees during this time period). (D) Weather data of temperature and sunlight intensity for local weather station MeteoBlue-Litzelstetten (2.2km from hive), plotted over time using the same time range as A-C.

The increase in drone speed was first detectable at age 5-6 days, became pronounced by age 7 days, and continued throughout the drone’s life (Figure 1A). The timing of the hyperactive bouts corresponded to when drones moved near to the entrance (Figure 1C) and spend time outside of the hive (Figure 1B), and so is likely associated with the onset of orientation/mating flights. These bouts tended to occur in afternoons, when outside temperatures were warm and the sun was out (Figure 1D).

Drones also changed their position in the nest as they aged. Young drones (ca. <10 days old) were located in the center of the nest, on comb containing brood (Figures 2, S1). From ca. age 10 days, drones shifted from the center of the nest to the periphery, i.e. the edge areas of each frame in the observation hive (Figure S3), which corresponded to areas bordering the brood. As they aged further (age 14-19 days), drones shifted closer to the nest entrance (lowest frame - see Figure 2); however, their average distance from the exit changed during the day (Figure 1C). After drone cohort 2 had reached age 10 (June 22), the two drone cohorts used similar areas of the nest. The shift from upper/periphery areas to the lower frame is noticeable for both cohorts on June 26 (age 19 and 14 days for drone cohorts 1 and 2, respectively - see Figure S1). This suggests that spatial positioning of drones in the nest is influenced not only by age, but also by the positioning of other drones.

**Figure 2:**
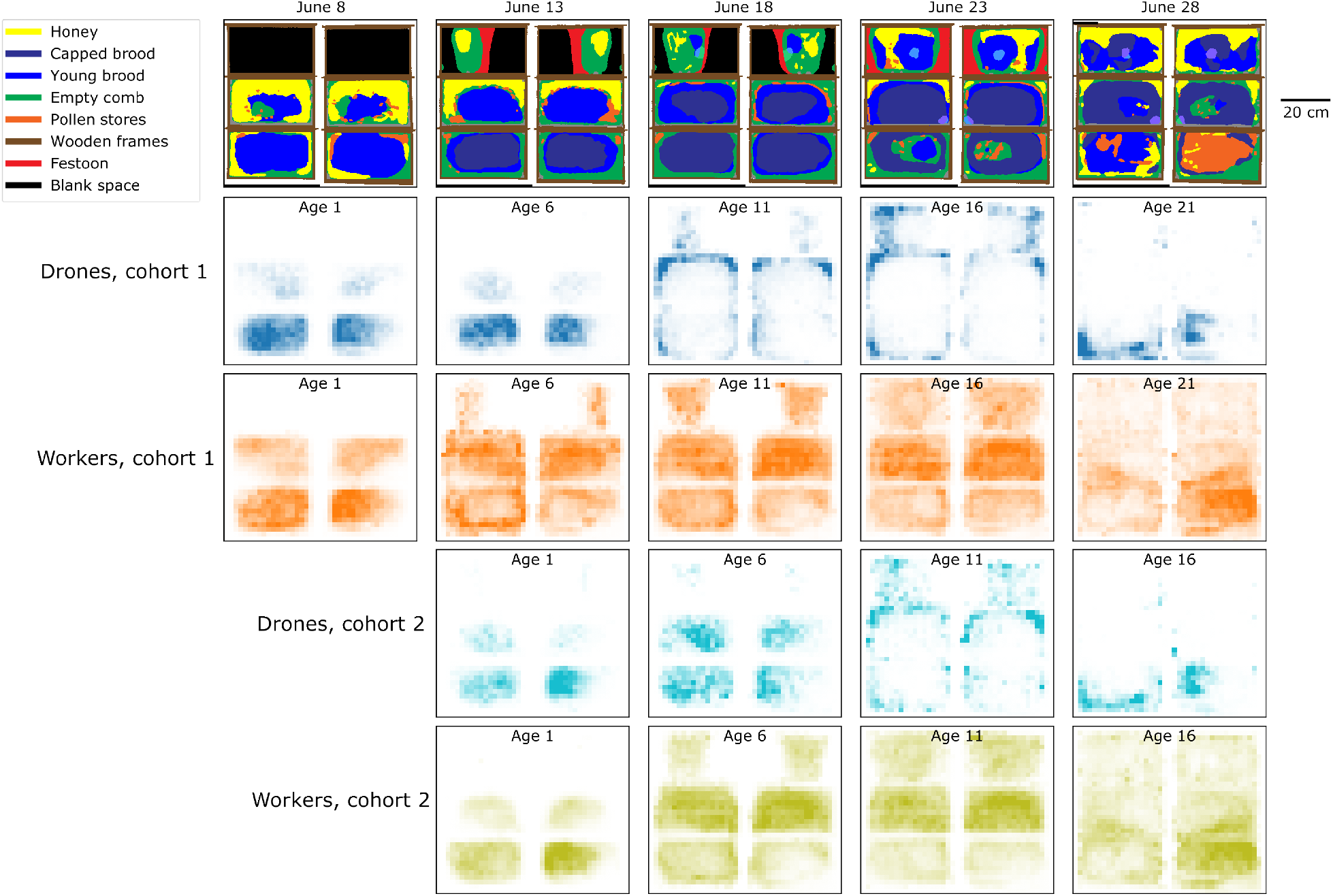
Spatial location of drones and workers. Nest contents over the observation period, and two-dimensional spatial histograms showing the locations of workers and drones each day, at 5-day intervals (see Figure S1 for all days). For each image, we display position in the observation hive by showing the back side on left, and the front side on the right; in this image, the nest exit is on the lower right corner (see also Figure S2)

During the observation period, the colony was also in the process of forming a festoon to build comb on the upper frame (see red ‘Festoon’ areas in Figures 2 and S1). June 26 was the last day the festoon was observed. After this, there was a notable increase in honey stores in the nest; honey areas went from 10.1% of the nest on June 26, to 12.1% on June 28, and then to 17.2% on July 2. During this time of increasing honey stores, all worker bees increased their time spent outside (fraction of all workers outside the hive after June 26; Figure 1B).

### 4.2 Synchronized activation of drones

We observed synchronization in drone activity, such that many drones became active at the same time—we refer to this as a “shared activation period”. We use the data to form a working definition of this period in order to enable quantitative analysis. To do this, we denote an individual drone as “activated” in a time bin *t* if its median speed is greater than a threshold of *s*_*act*_, or if the drone is identified as being outside of the nest during *t*. We use a high value of median speed for the threshold: *s*_*act*_ = 0.5 cm/sec, which corresponds to the 92.8% quantile of median speed values for all tracked bees in the observation period of June 7-29. We define activation using high speed activity OR outside the nest because these activities coincided, because in-nest speed in-between flights tended to be high, and because these are mutually exclusive activities(i.e. in-nest speed does not exist when drones are outside the nest). To compare the onset and duration of activity on different days, we then define the “shared activation period” as when >25% of drones age >=7 days are activated at the same time (Figure 3A). The age threshold of >=7 days old is used because drone activation was not strongly observed until this age. We again note that the definition of the shared activation period is made in order to enable quantitative analysis; the parameters are chosen such that this period is a meaningful representation of the data (Figure 3A), and the definition is not overly sensitive to the exact parameter values for speed threshold and fraction of drones (Figure S4).

**Figure 3:**
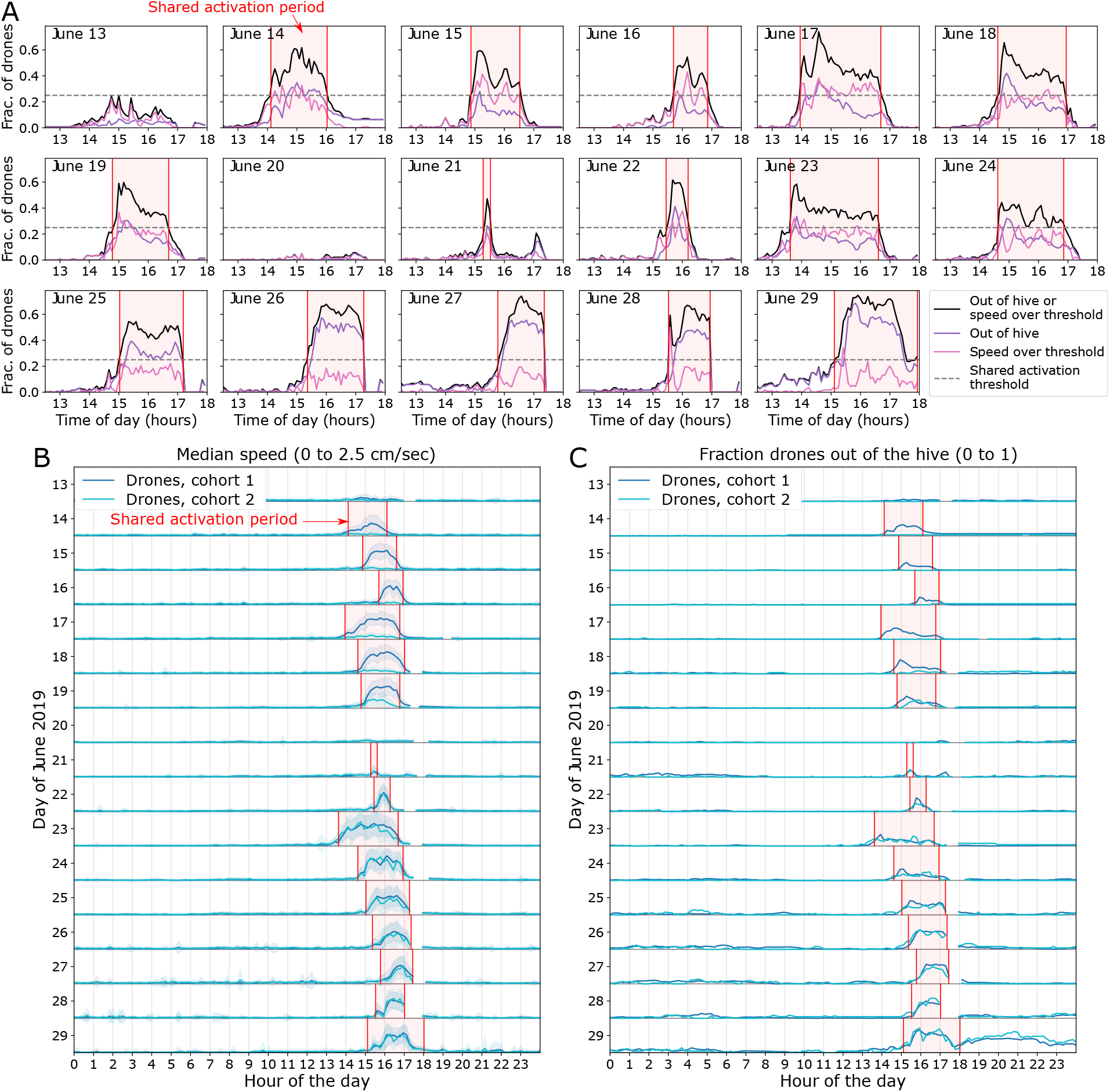
Shared activation period and hourly activity. (A) We define the “shared activation period” to begin when >25% of drones age >=7 days have high speed (greater a threshold of *s*_*act*_ = 0.5cm/sec) or are outside of the hive, and to end the last time when <25% of drones satisfy either condition. Starting from June 14, every day had a shared activation period, with the exception of June 20. Note that on June 25-28, we were able to detect the onset of the shared activation period, but the end of the period was obscured by the comb contents measurements. See also Figure S4 for the sensitivity of the collective activation period to the parameters of fraction of drones and *s*_*act*_. (B) Median speed, averaged across each cohort (y-axis range for each day is 0 to 2.5 cm/sec) and (C) Fraction of drones out of the hive, for each cohort (y-axis range of 0 to 1 for each day). Each day is aligned by hour (x-axis). Gaps in the data shown are due to the comb contents measuring period, which requires the cameras to be temporarily paused.

After a shared activation period was first observed on June 14, all subsequent days June 14-29 had a shared activation period with the exception of June 20. Although the shared activation period varied from day to day, it always began between 13:00 and 16:00, and always ended by 18:00. On some days, as many as 70% of the drones were active during this period. The duration of the shared activation period varied from 3 hours (June 23) to only 15 minutes (June 21) (Figure 3).

Individuals in drone cohort 2 started from age 7 (June 19) to increase their speed and take part in the shared activation period, becoming activated at the same time as drone cohort 1 each day (Figure 3B-C). Therefore, even though these different drone cohorts were of different ages, they were synchronized in activity such that we observe only a single shared activation period. During the same time periods, worker bees did not exhibit a particular increase in activity levels (Figures S6, S7).

The duration and onset of the shared activation period depended on the daily weather conditions. Of the quantities examined, sunshine duration had the highest positive correlation with duration of the shared activation period (Figure 4). On one day (June 20), we did not observe a shared activation period; the drones remained immobile throughout the day. Using local weather data, this day had fewer hours of sunlight (<5 hours sunlight recorded at both nearby weather stations DWD-Konstanz (1.6km from hive) and MeteoBlue-Litzelstetten (2.2km from hive) - see Figure S8). On the following day (June 21), some drones did increase their speed and took trips outside (over 40% of drones became activated for a brief amount of time on this day – see Figure 3A), but this day had the shortest activation period (15 minutes). On each day, the onset of the shared activation period occurred slightly after the period of strongest sunlight (Figure S9).

**Figure 4:**
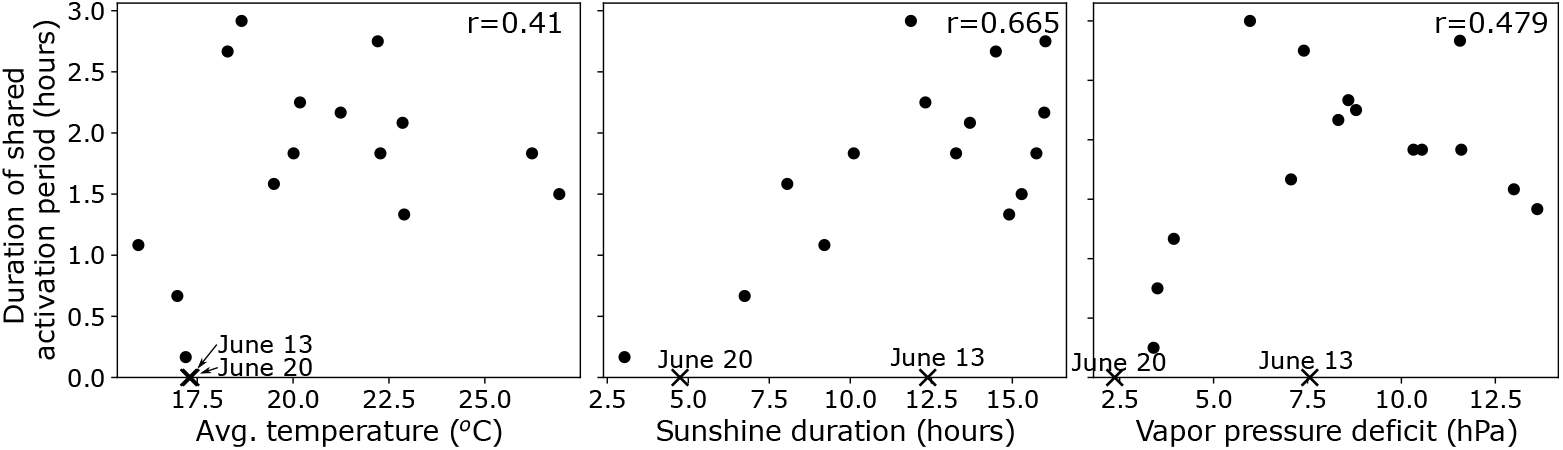
Duration of shared activation period and daily weather conditions. Duration of the shared activation period (y-axis) is plotted as a function of daily weather averages or totals (x-axis). Shown are average daily temperature, total sunshine duration, and average vapor pressure deficit obtained from MeteoBlue weather station Konstanz-Litzelstetten. See also Figures S8, S9 for more details and other weather measurements during the observation period, including additionally precipitation, pressure, and average wind speed. June 13 data point is labeled to show the date before drones became activated; June 20 is labeled to show the date that did not have a shared activation period. The correlation values (*r*) are shown in the upper right of each plot. Using a p-value threshold of 0.05, only the correlation with total sunshine duration is significant (p values of 0.102, 0.004, and 0.052 for temperature, sunshine duration, and vapor pressure deficit, respectively).

### 4.3 Individual drone activity onset

We next examine the activity patterns of individual drones. We see remarkable synchronization in the onset and end of hyperactivity and outside trips among the individual drones (Figure 5). During active periods, drones exited the nest multiple times, and maintained high movement speed when they returned to the nest. The number of outside trips per drone during the shared activation period was 1.17 ± 1.05, meaning that most drones took 1-2 trips (although some drones did exit many more times - see Figure S5A). The average time per outside trip during the activation period was 24.2 ± 11.6 minutes. Drones tended to increase their speed before exiting the nest (Figure S5B). The end of the activation period saw individuals returning into the nest and reducing their speed, showing that not only the onset of activity, but also the decrease, is synchronized among drones (Figure 5).

**Figure 5:**
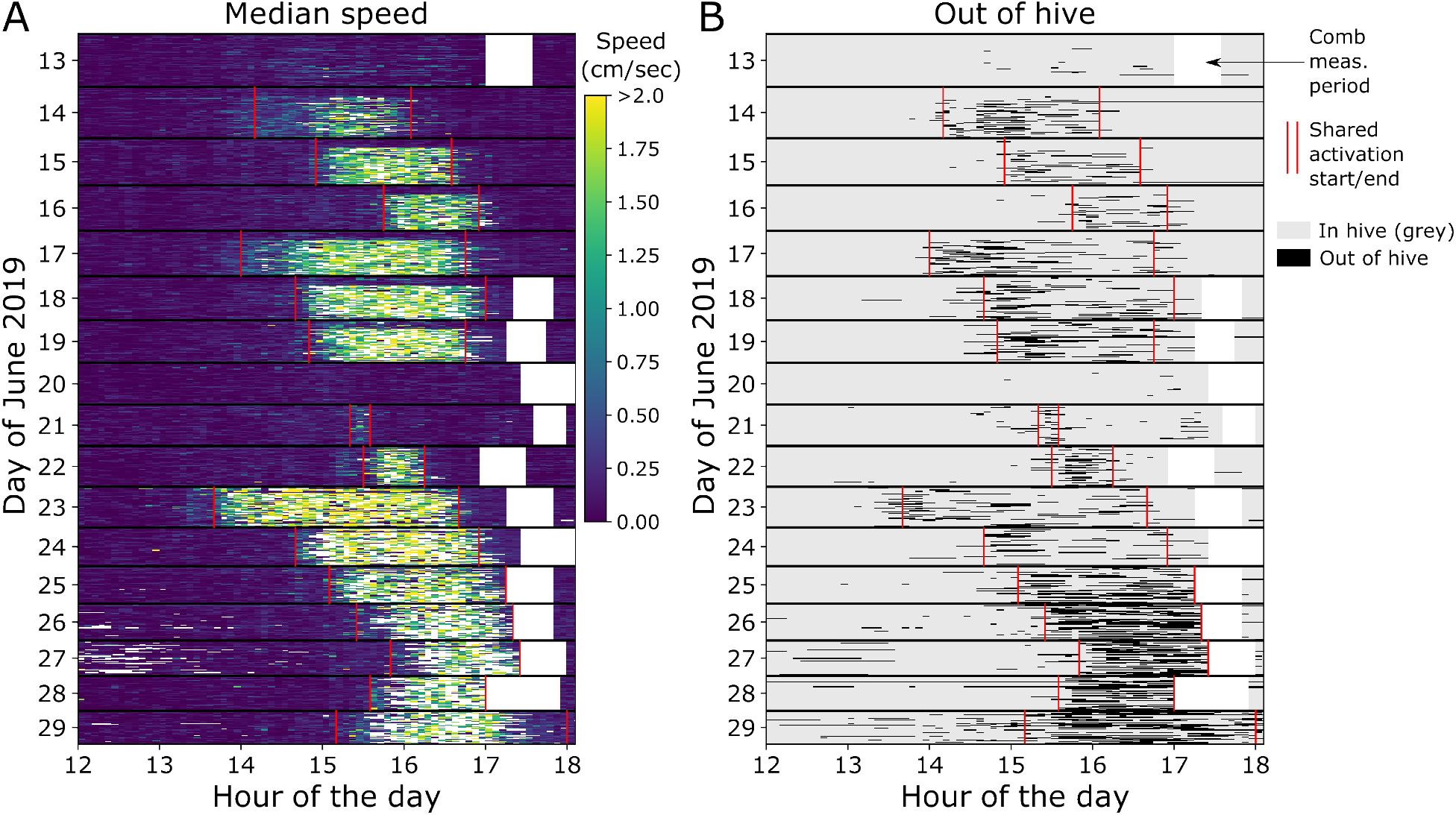
Individual activity patterns. Raster plots, where each row represents an individual drone on a particular day, showing (A) Speed and (B) Trips out of the hive, in 5-minute bins (see Methods for data processing details). White areas are when cameras were temporarily blocked to transcribe nest contents, and red lines denote the start/end of the shared activation period (see also Figure 3). On June 25-28, we were able to detect the onset of the shared activation period, but the end of the period was obscured by the comb contents measurements. Figure S7 for analogous plots with worker cohorts 1 and 2.

To determine if the drones were consistent in their onset of activity (e.g. drones that were consistently “early-to-activate” or “late-to-depart”), we defined the individual activation time for drone *i* as the first time bin *t* starting from an hour preceding the shared activation period where either its speed *s*_*i*_ *> s*_*act*_, or the drone was out of the hive. We then calculate the correlation in activation timing of all drones from one day to the next. The correlation was nonzero and varied across days, but was generally low (Figure 6A). This suggests that there is not a strong tendency for certain drones to consistently be first or last to become activated, or for certain drones to consistently “lead” the onset of shared activation each day.

**Figure 6:**
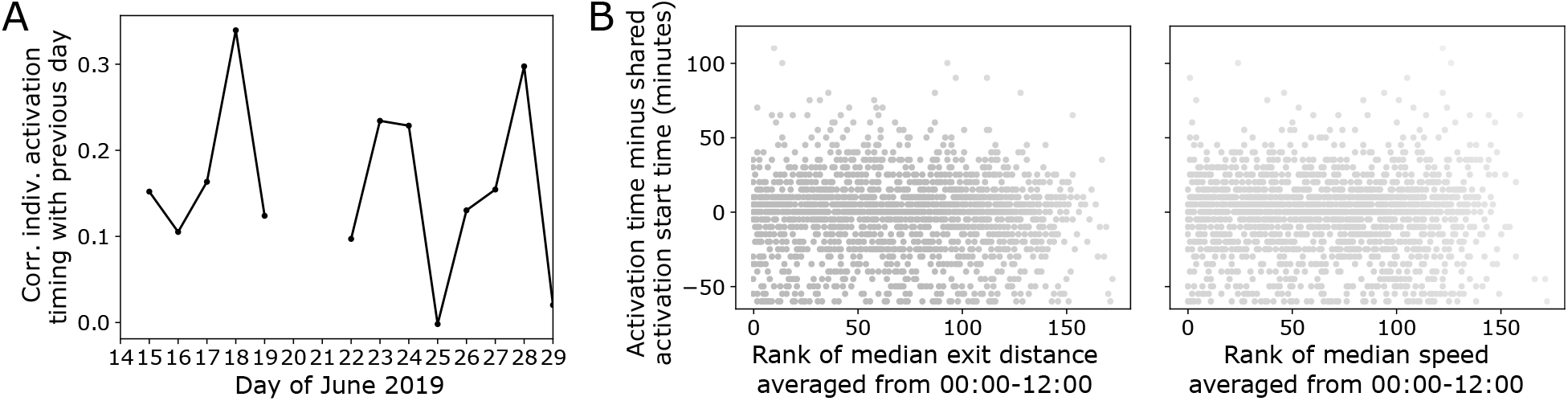
Testing for consistency in activity onset. (A) Correlation of individual activation time from one day to the next. The value for a designated day is the Pearson correlation of the timing of drones that became activated on that day, with the timing of drones that had also become activated on the previous day. Because there was no shared activation period observed on June 20, the correlation is not calculated for June 20 and 21. (B) Scatter plot showing ranked average morning-time (00:00-12:00) median exit distance (left) or speed (right) versus individual activation time. All days with shared activation periods are included; each point represents one drone on a single day. The ranked value of exit distance or speed is with respect to other drones on that day; this is done because drone positioning changes over developmental time (See Figure 2). The overall correlations are low: the correlation of ranked exit distance with individual activation time is -0.051, and of ranked speed with individual activation time is -0.017.

We further asked if previous behavior or location predicts activation timing of individuals with respect to the shared activation period. We used the morning hours of 00:00-12:00 each day to calculate average values of exit distance and speed for each drone. Because drones change their location in the nest over time (see Figure 2), we used a ranking of drones within each day. Figure 6B shows that there is nearly no correlation of either morning-period exit distance, or speed, with individual activation timing.

### 4.4 Model of social activation of drones

The synchronization of the activation period (Figure 5) suggests that social information exchange, in addition to external weather-related factors, drives individual activation. To consider this mechanism in detail, we formulate a simple threshold model and simulate the response of a group of agents with and without social information exchange. The purpose of this model is not to specifically match the observed results, but rather to determine what conditions would be necessary to observe the same general behavioral patterns (e.g. presence/absence of social information, individual thresholds, noise levels). For other examples of threshold models applied to social insects see Beshers and Fewell (2001); Jandt and Dornhaus (2014); Ulrich et al. (2021). In the model, an individual agent *i* becomes activated when its internal decision state *x*_*i*_ reaches a threshold of *η*_*i*_. Each of the *N* agents in the group can have a different threshold value. An individual’s decision state is described by an Ornstein-Uhlenbeck process (Smith, 2000), as a leaky accumulator of social information and an external signal, plus noise:

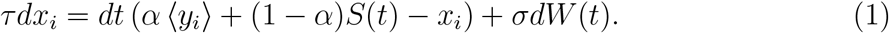

In this equation, *τ* is the timescale for changes in the decision variable, *σ* is the amplitude of the noise in the decision variable, and *dW* (*t*) is a Wiener noise process. The parameter *α* sets the weighting of social information versus the external signal. Social information is represented by the fraction of group members that have already crossed their decision thresholds, i.e ⟨*y*_*i*_⟩, where ⟨. ⟨ represents an average, and *y*_*i*_ = 1 if *x*_*i*_ ≥ *η*_*i*_ and otherwise 0. The external signal is set to *S*(*t*) = *A* sin(2*πt*) in order to represent periodic changes such as daily rhythms or sunlight, and the amplitude *A* sets the strength of the signal. See Figure 7A for an illustration of the model, showing the external signal and decision states of 3 simulated agents.

**Figure 7:**
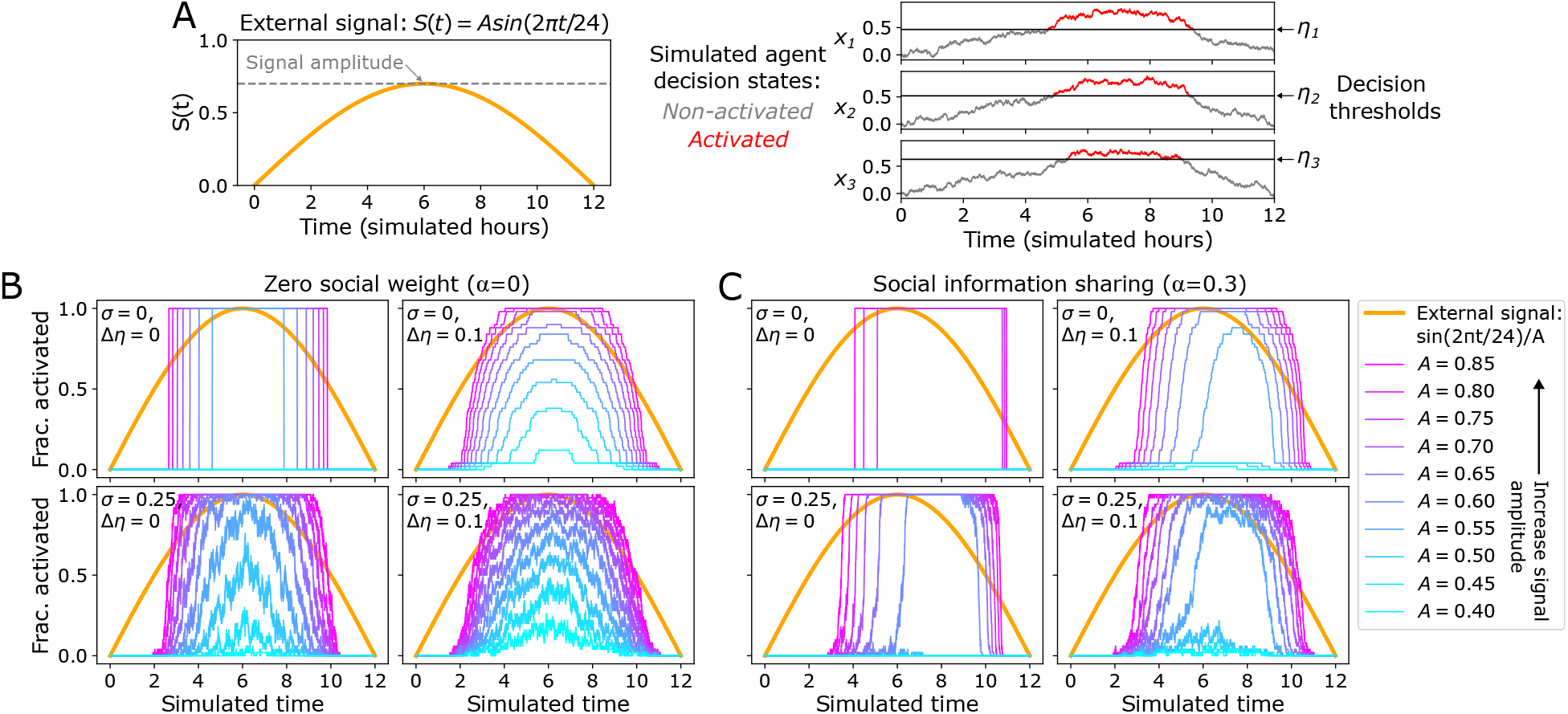
Activation model with external and social information. We model the activation of drones using an accumulation-to-threshold model, where evidence comes from both an external signal as well as social information (social information is the fraction of group members already activated). (A) Model illustration. The external signal is sinusoidal to represent circadian daily patterns, with amplitude set by the parameter *A*. Individual agent decision states *x*_*i*_ follow a leaky accumulator process with noise amplitude *σ* (Eq. 1). Across a simulated group of *N* individuals, each agent’s decision threshold *η*_*i*_ is drawn from a normal distribution with standard deviation *δη*. The parameters *α* sets the fraction of evidence due to social information: *α* = 0 is only external signal, and e.g. *α* = 0.3 represents 30% social evidence and 70% external signal used by each agent. (B-C) Simulation results for zero vs nonzero values of decision noise (*σ*) and within-group threshold differences (∆*η*), shown for (B) zero social information use, and (C) both social information and external signal used as actuation evidence. In each plot, shown is the normalized external signal, and results for fraction of group members activated over time in response to different signal amplitudes *A*. Each 2 × 2 grid shows results for (top row) zero decision noise, (bottom row) nonzero decision noise, (left column) equal thresholds, and (right column) threshold differences among group members. Simulations use a group of *N* = 50 agents.

We simulate different parameter settings for noise, threshold differences among group members, and social information use to ask what mechanisms are consistent with the main observations in the data. The parameter *σ* represents noise in an individual’s decision process (“decision noise”). We set individual agent threshold values using the distribution *η*_*i*_ =𝒩(0.5, ∆*η*), such that ∆*η* represents “threshold differences” among individuals. For different combinations of decision noise and threshold differences, we compare simulations of zero versus nonzero social information exchange (as set by the parameter *α*).

In the data, we see that on days with more sunshine, the shared activation period is longer (Figure 4). The signal amplitude *A* can be considered to represent the strength of the sun in the day (orange parabola in Figure 7); for all parameter values, the model shows the trend of increasing activation time with higher *A*. We see, however, that the synchronization and time of onset of the activation period depends on the values of noise, threshold differences, and social information weight (Figure 7B,C). With zero decision noise and all agents having the same threshold (*σ* = 0, ∆*η* = 0), we see perfect synchronization among agents, regardless of social information use: all group members are either activated or not activated (top left subplot in Figure 7B,C). A more realistic case considers noise in the decision-making process to represent individuals as imperfect sensors of their environment (*σ* > 0, bottom row of Figure 7B,C). If either decision noise or threshold differences are present, then the overall fraction of the group activated shows gradual changes with signal amplitude, i.e., no longer a synchronized all-or-none response (Figure 7B).

The use of social information increases the synchronization of activation among group members. Even with decision noise and threshold differences, a group with social information sharing can synchronize the activation response (Figure 7C). The use of social information also speeds up the onset of the activation period (how long it takes from the first activated individuals until approximately all are activated), as well as shifting the time of activation onset to relatively later (c.f. Figures 7B and C). Therefore, agents that incorporate social information give results that more closely align with our observed data than agents that do not incorporate social information.

## 5 Discussion

Using long-term automated tracking, we followed the in-nest behavior of honey bee drones, examining how they move and where they position themselves in the nest over their entire lives. While drones did spend the majority of their time immobile in the nest, we also found that drones are synchronously hyperactive inside the nest during afternoon periods that coincide with trips outside (Figure 1). This behavior developed with age, becoming prominent from age 7 days onward, in both drone cohorts. We defined the “shared activation period” as the time when a large fraction of drones were active – i.e. fast-moving or taking trips outside (see Figure 3) – and found that the onset occurs in the afternoon just after the sun is at its peak, and that the duration is longer on days with more sunshine hours (Figures 4, S9). Looking at individual drones, the onset and end of activation was synchronized – drones became active and started/ended their outside trips at the same time as other drones (Figure 5). However, individual drones did not show a strong consistency in their relative activation timing from day to day – i.e., there are not consistent “early-to-activate” or “late-to-depart” drones (Figure 6A). We simulated a threshold-based model of drone activation to demonstrate that the main observations in the data are consistent with individual decisions using both external and social information to decide when to activate (Figure 7).

Daily changes in activity can be triggered by external factors (such as weather), internal factors (such as circadian biological rhythms), or by social information exchange with other individuals. It is well established that honey bee workers exhibit circadian rhythms, which help organize roles within the colony, and can be socially mediated (Medugorac and Lindauer, 1967; Bloch and Robinson, 2001; Eban-rothschild and Bloch, 2012). This likely also applies to the timing of drone departure flights. We saw that the onset of the shared activation period occurs shortly after the time of strongest sunlight (Figure S9), and that the duration of the shared activation period is correlated with the weather conditions for that day (Figure 4C). While we did observe a positive correlation between the duration of the shared activation period and daily sunshine hours 4), is it possible that this trend could become saturated, or even reversed, under more extreme environmental conditions (e.g. less activation if the temperature is “too hot”). Given that we did not perform manipulative experiments to alter daily light exposure, we cannot directly distinguish between external cues from the sun, and internal triggers from biological rhythms. However, because biological rhythms take multiple days to establish or change (Roenneberg et al., 2003), and weather is highly stochastic, the absence of an increase in activity on the poor weather day (June 20) suggests that external factors, like weather, may play a stronger role than internal factors.

Specific experimental manipulations would be needed to test the relative importance of internal, external, and social factors for the timing of drone hyperactivity. There is suggestive evidence that internal factors determine flight time in drones; researchers used a flight room to clock-shift drones, and found that when moved outdoors, the drones retained their clock-shifted departure time (Pfannenstiel and Koeniger, 2000). These findings, however, were published as an abstract, and so would need to be confirmed. Ideally, such an experiment would also investigate the in-nest hyperactivity that we observed, to potentially determine whether clock-shifted drones could socially-induce other non-shifted drones into a hyperactive state.

The timing of individual activation did not correlate with morning-time spatial position or speed of individual drones (Figure 6B). Conspicuous hyperactivity may itself be the social cue that helps to synchronize all the male reproductive drones across the colony, but the precise mechanisms for information transmission and activation remain to be explicitly tested. This could include the potential for chemical communication; drones have been shown to exhibit age-based attraction to conspecifics (Bastin et al., 2017). In our model of drone activation, we did not consider the mechanisms of information transmission, but instead assumed that individual drones can sense when others are activated (Eq. 1). An interesting avenue for future work is to examine the fine-scale dynamics of how drones interact and respond to other drones, including social communication mechanisms using interactions and proximity among drones (Wild et al., 2021). In conjunction with specific experimental manipulations, one could determine how internal, external, and social factors influence activation. Testing how drones coordinate their in-nest behavior may also provide additional insights into the potential for coordination outside of the nest (reviewed in Mariette et al. (2021).

Our model of drone social activation represents individuals as leaky integrators of an external signal and social information. Using social information can reduce uncertainty or noise in estimating environmental states or making decisions (Srivastava and Leonard, 2014; Ellison et al., 2016). We see similar results in our simulations: Figure 7 shows that synchronized activation can occur if independent agents have zero noise and identical activation thresholds, or if social information is used in the presence of noise and threshold differences. Accumulation-to-threshold models have been applied to many different systems, with applications including individual neuron spiking (Teeter et al., 2018), perceptual decision making (Usher and McClelland, 2001), foraging (Davidson and El Hady, 2019; Bidari et al., 2022), and economic or purchasing decisions (Gluth et al., 2012; Krajbich et al., 2012). We used a leaky accumulator model formulation, which previously has been applied to tasks that require a detection of signal changes (Clifford and Ibbotson, 2002; Glaze et al., 2015). While we represented social information sharing with a simple fraction of activated drones, we note that future work could incorporate additional mechanisms of information sharing (e.g. local interactions among neighbors or nonlinear forms of coupling (Zhong and Leonard, 2019; Bizyaeva et al., 2021).

The onset of in-nest hyperactivity aligns with important stages in drone sexual maturation (reviewed in Koeniger et al. (2014); Rangel and Fisher (2019)). Drone departure flights first occur between age 6-9 days old (Reyes et al., 2019); sperm move from the testes to the seminal vesicles between 7-8 days (Snodgrass, 1956; Mackensen, 1955); and sperm viability peaks as early as 7 days old (Locke and Peng, 1993). While these physiological changes coincide with the behavioral changes we observed in the nest, it remains unknown the extent to which these changes are coupled.

Drones and virgin queens have a common interest in meeting in the most efficient way possible. There are two mechanisms by which independent parties can indirectly coordinate meetings without direct communication: (1) restrict the spatial component, and (2) restrict the temporal component (Fenster et al., 1995; Dostálková and Špinka, 2007; Couzin, 2018). Drone congregation areas were already known to restrict the spatial component, and afternoon flights restrict the temporal component. Here we show that the restricted temporal component is also associated with a limited period of drone hyperactivity *inside* the nest. Although more time spent outside is coupled with higher mortality (Visscher and Dukas, 1997), drones are also under selective pressure to be present in the drone congregation area *before* virgin queens arrive – if they have not arrived in time, they will miss their opportunity to mate. These two forces contrast with one another: the need to arrive early to overlap with virgin queens is a pressure to extend time outside, but mortality risk limits the time spent outside. Combined, these factors appear to be a form of stabilizing selection on the drone’s behavioral phenotype (Hansen, 1997), which results in a strong “on/off” activation period. In locations with high mortality outside of the nest, such as where the bee-eating bird (*Meropidae*) is common (Ali and Taha, 2012; Loope, 2015), we would therefore predict a shorter period of drone activation than in locations where mortality is lower (Smith, 2018). In places with high outside-nest mortality, we would still expect to see synchronized hyperactivity, but over a shorter period. While our study did observe in-nest hyperactivity across all tagged drones, it would also be interesting to see how these patterns vary across colonies and environmental conditions.

A honey bee colony is an integrated superorganism, with workers functioning as soma, and drones as gametes (Seeley, 1989; Smith and Szathmáry, 1995). Just as the optic nerve and the testes serve different roles within a multicellular organism, so too do the individual bees that form a colony. It is unsurprising that drones are referred to as “lazy” (Frisch, 1954; Gadagkar, 2021) - they do indeed spend the majority of their time immobile at the periphery of the nest. However, this behavior can also be viewed as adaptive: by remaining immobile they conserve their energy until mating time, and by moving to the nest periphery, drones prevent potential obstruction to workers in action. Just as drones have morphological and physiological adaptations to maximize their mating opportunities (e.g. large eyes (Menzel et al., 1991); no hypopharyngeal glands (Hrassnigg and Crailsheim, 2005)), their behavior is also adaptive, both inside and outside of the nest. Using long-term automated tracking, we found that drones, in contrast to their reputation, can be *the most active individuals in the colony*, albeit for a limited time each day. Therefore, drones do exhibit specialized in-nest behavior, which highlights their role as the male gametes of the colony.

## 6 Acknowledgements

Thank you to Giovanni Galizia for providing access to his rooftop apiary, to Jan Peters for expert colony maintenance, to Dagmar Olalere and Katja Anderson for essential logistical support, to Markus Miller for IT expertise, and to Sinje Tigges, Christine Bauer, and Jayme Weglarski for patiently helping to tag honey bee workers and drones.

## 8 Data and code availability

All data is available though Zenodo (doi.org/10.5281/zenodo.7298798), and code to reproduce the results in the paper is available on Github (jacobdavidson/bees_drones_2019data).

---

**Figure S1:**
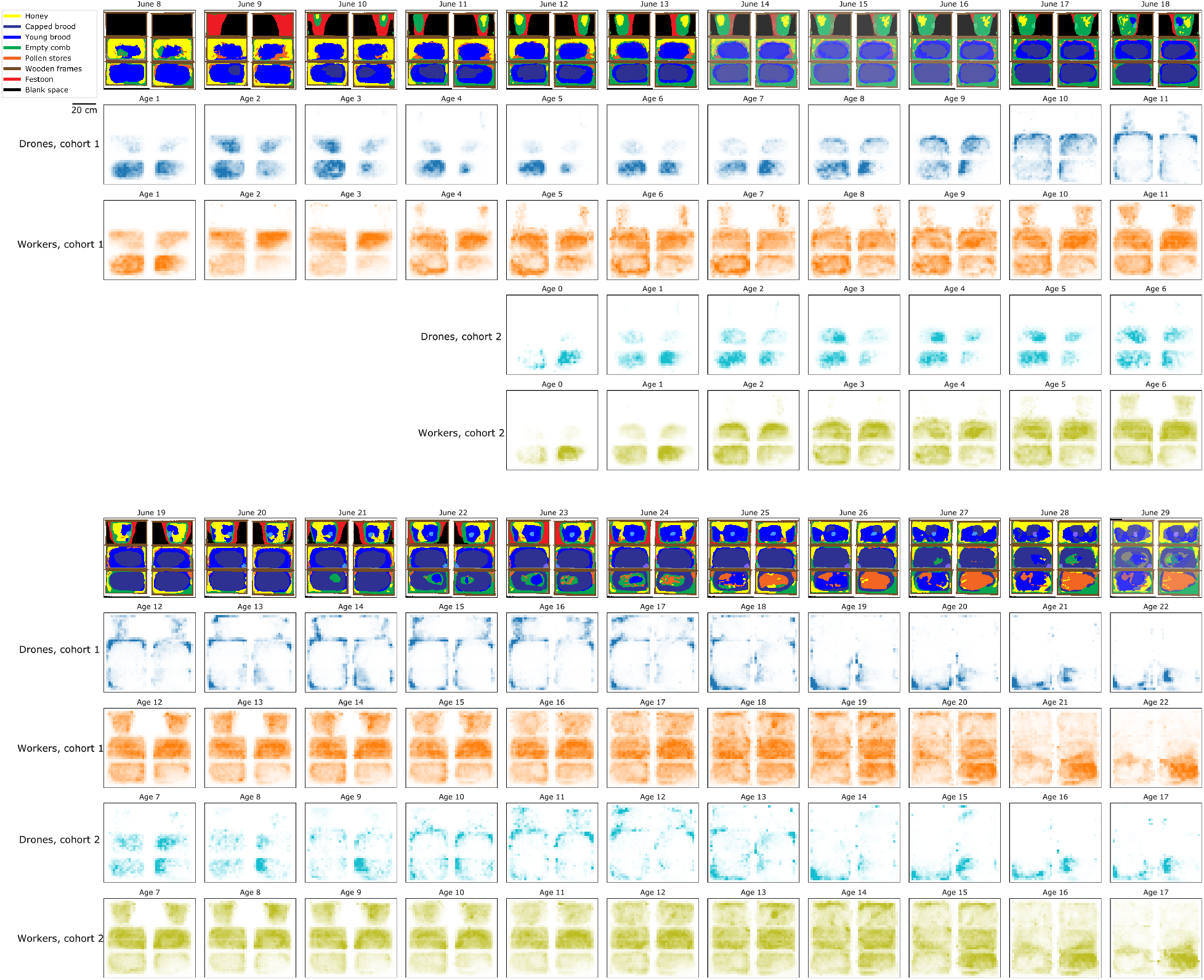
Two-dimensional spatial histograms for each day with age. Analogous results to those shows in Figure 1B, but including all observation days. The top row shows nest contents over the observation period, and two-dimensional spatial histograms show the locations of workers and drones on each day. For the days where comb measurements were not taken (June 14, 15, 16, and 29), the comb is not known exactly, so both of the nearest measurement days are shown overlayed, with the transparency weighted proportionally to the time from the measurement day (i.e. less transparency for the closer measurement day).

**Figure S2:**
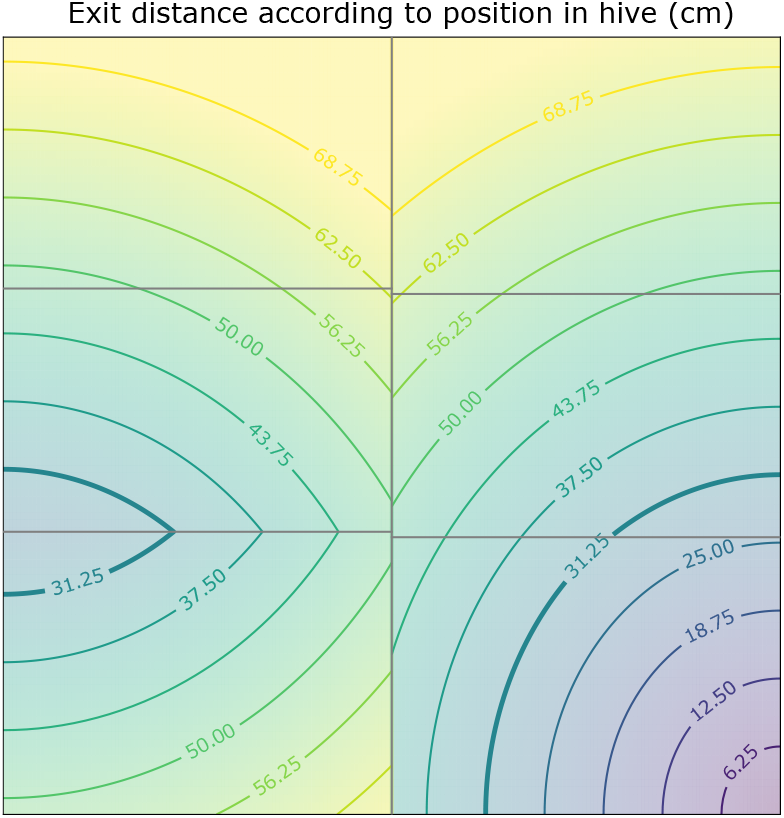
Exit distance. Exit distance for different locations in the observation hive. Note that the exit is located at the lower right of the front side of the observation hive (lower right of images), and that crossings from one side to the other are possible on the frame borders. In order to estimate when bees exited the nest, we used a threshold of 31.25cm on median exit distance in the 5-minute bins (see Methods for details on this algorithm); this value is made bold in the figure.

**Figure S3:**
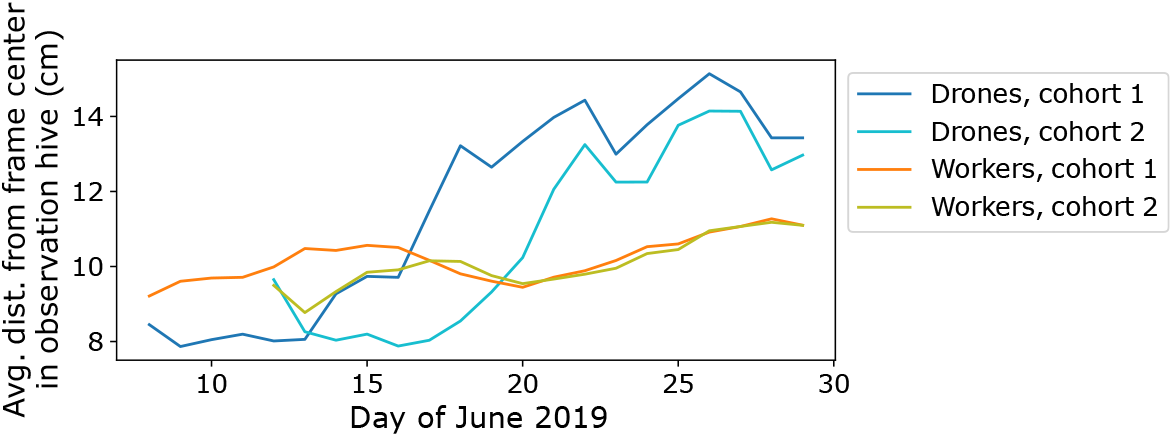
Average distance from frame center. The observation hive consists of 6 frames, with 3 on each side (see Figures 2, S1). Shown is the average distance from center of the current frame, comparing drone versus worker over time.

**Figure S4:**
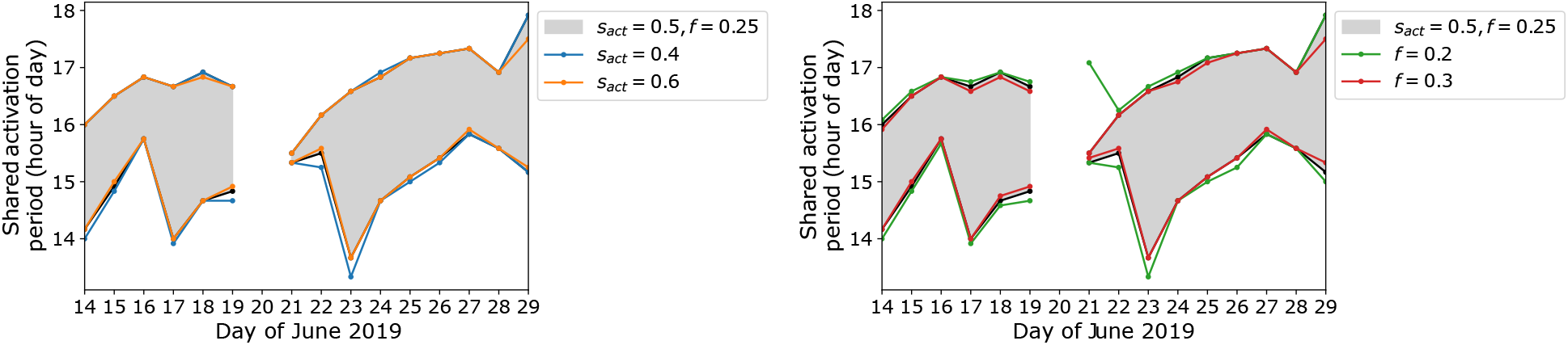
Sensitivity of shared activation period definition to parameter values. In the main analysis, the value of speed threshold used was *s*_*act*_ = 0.5, and the fraction of drones *f* = 0.25. Shown is how the duration of the shared activation period depends on (left) changing the speed threshold *s*_*act*_, and (right) changing the fraction of drones threshold (*f*). While the definition is not sensitive to the speed threshold, individual days have patterns that are sensitive to the value of *f* - in particular, using a lower drone fraction threshold defines a longer shared activation period for June 21, because it then would extend to include the whole time until the small ‘spike’ in activity near 17:00, which follows a period of low speed / inside. The parameters of *s*_*act*_ = 0.5 and *f* = 0.25 used in the main analysis were chosen by visual inspection, to be representative of capturing the observed quick start and end of the shared activation period (see Figure 3A).

**Figure S5:**
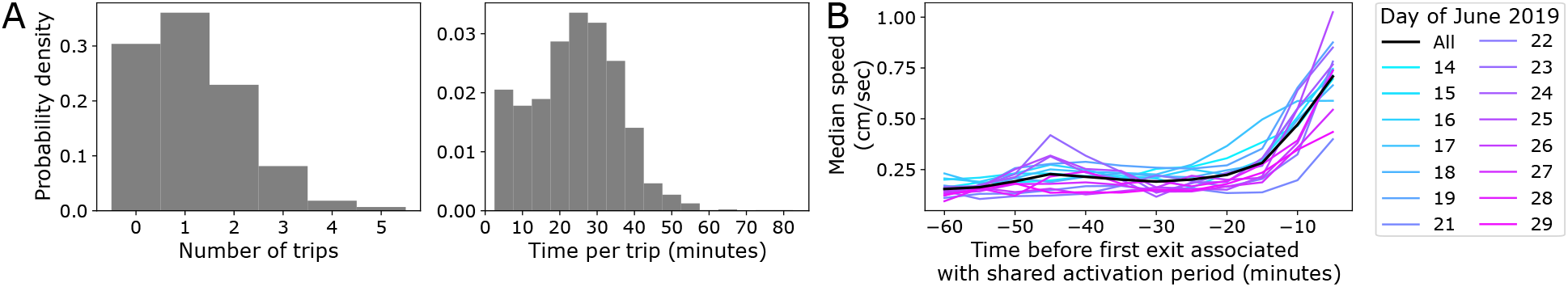
Individual drone outside trips and speed before exiting. (A) The distributions of number of trips per drone (left) and time per trip (right), for drones during the shared activation periods. Distributions include data across all days with a shared activation period. (B) Median speed of drones, in the time period preceding their first exit associated with the shared activation period (first exit time defined as the first time bin *t* starting from an hour preceding the shared activation period where the drone was out of the hive).

**Figure S6:**
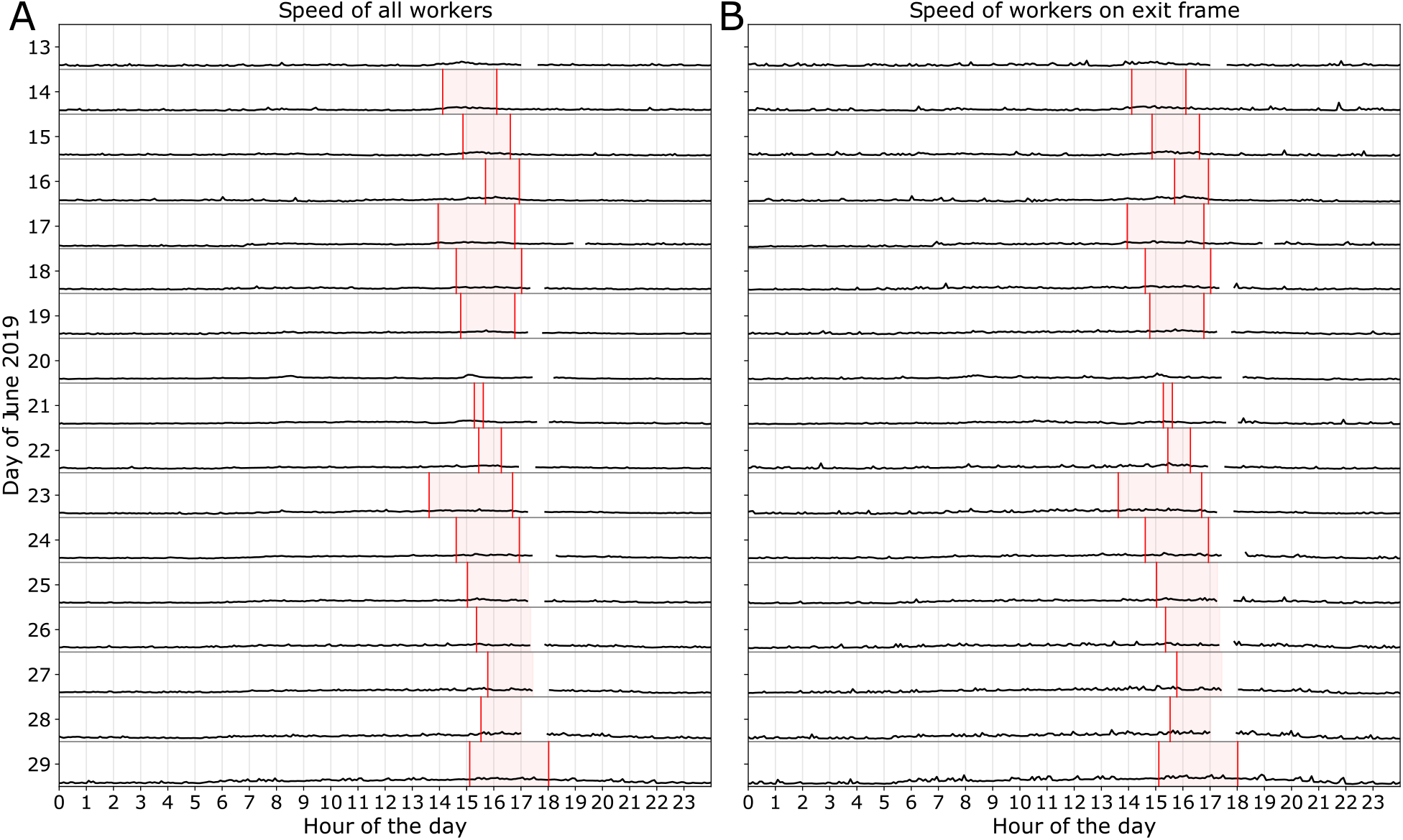
Average movement of workers during each day. The average speed of workers over time each day, calculated as the mean of the median 5-minute-bin speed of workers. The data is plotted analogous to Figure 3B, with the shared activation period of the drones highlighted in red, and range for each day is 0 to 2.5 cm/sec (this is the same scale as Figure 3B). Shown are averages for (A) all workers, and (B) workers whose fraction of time observed on the exit frame in a given 5-minute time bin was at least 50%.

**Figure S7:**
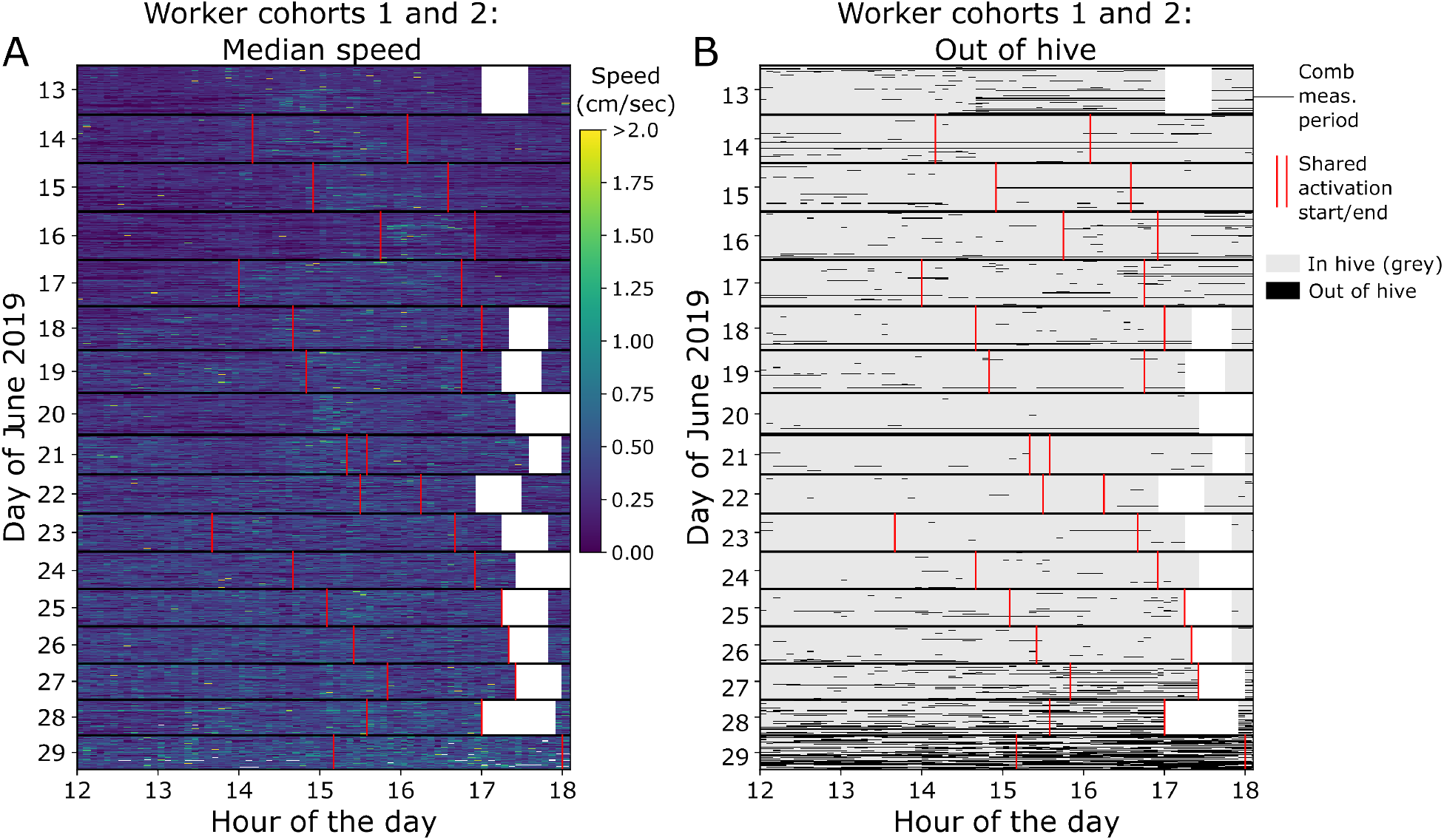
Worker bee cohorts: individual activity patterns. Analogous plots to Figure 5, but showing worker bee cohorts 1 and 2 (who have the same birth dates, respectively, as drone cohorts 1 and 2). In the raster plots, each row represents an individual bee on a particular day, showing (A) Speed and (B) Trips out of the hive, in 5-minute bins (see Methods for data processing details). As in Figure 5, white areas are when cameras were temporarily blocked to transcribe nest contents, and red lines denote the start/end of the shared activation period of the drones (see also Figure 3B-C).

**Figure S8:**
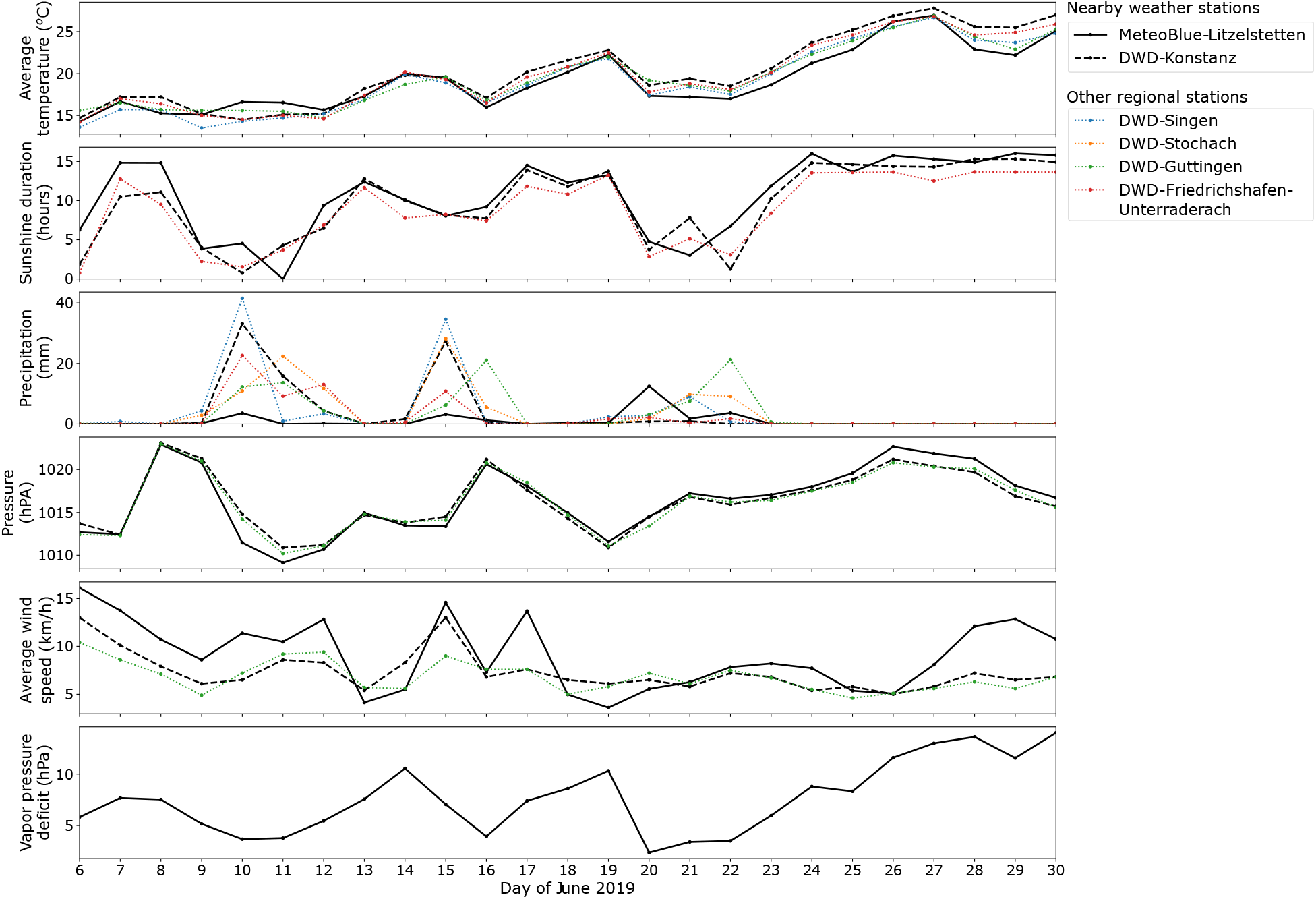
Daily weather for observation days. Nearby and regional weather data for observation days, showing average temperature, sunshine duration, total precipitation, average atmospheric pressure, average wind speed, and average vapor pressure deficit for each day. Not all measurements are available for all weather stations. Recorded values of precipitation and wind speed sometimes differ for the different weather stations, but other measurements are very similar. Nearby station Konstanz-Litzelstetten data is from MeteoBlue (2.2km from hive), and other data is open data from the Deutsche Wetterdienst (DWD). Station Konstanz is the closest at 1.6km from the hive.

**Figure S9:**
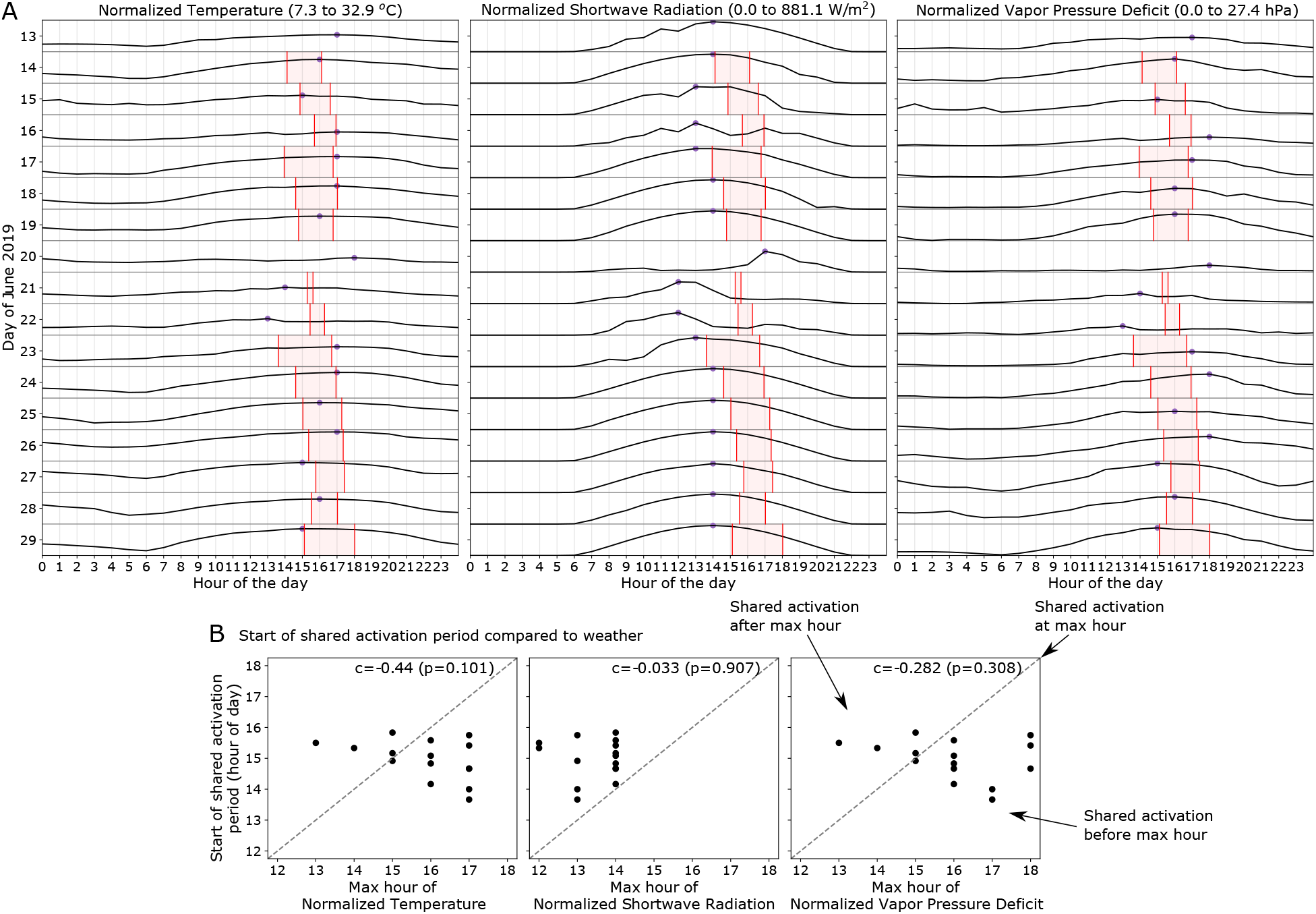
Hourly weather and start of activation period. (A) Detailed hourly temperature, sunshine intensity, and vapor pressure deficit recorded at Konstanz-Litzelstetten (MeteoBlue - 2.2km from the hive), for selected observation days. This data is plotted analogous to speed and number of drones out of the nest that are shown in Figure 3B-C, with the shared activation period highlighted in red. Points show the time of day for the maximum value of each quantity. (B) Start of the shared activation period (y-axis) plotted as a function of time of peak weather quantities (x-axis), using the peak times shown in A. Correlation and p-values are shown on each plot. The dashed lines shows the expected trend if the start of the shared activation period coincided with the maximum hour of each weather quantity. See also Figure 4 for a comparison of daily weather averages for nearby stations with the duration of the shared activation period.

Figure S10: **Supplementary Video.** Drone trajectories within the observation hive from 12:00 to 18:00, 14 June through 29 June 2019. Trajectory color denotes average speed. Individual trajectories are shown atop a grey background, with white denoting the comb measurement period (when cameras were briefly turned off). Note that 20 June (blue text) is the day during which there was no shared activation period.

